# Prenatal vitamin D deficiency alters immune cell proportions of young adult offspring through alteration of long-term stem cell fates

**DOI:** 10.1101/2023.09.11.557255

**Authors:** Koki Ueda, Shu Shien Chin, Noriko Sato, Miyu Nishikawa, Kaori Yasuda, Naoyuki Miyasaka, Betelehem Solomon Bera, Laurent Chorro, Reanna Doña-Termine, Wade R Koba, David Reynolds, Ulrich G. Steidl, Gregoire Lauvau, John M. Greally, Masako Suzuki

## Abstract

Vitamin D deficiency is a common deficiency worldwide, particularly among women of reproductive age. During pregnancy, it increases the risk of immune-related diseases in offspring later in life. However, exactly how the body remembers exposure to an adverse environment during development is poorly understood. Herein, we explore the effects of prenatal vitamin D deficiency on immune cell proportions in offspring using vitamin D deficient mice established by dietary manipulation. We found that prenatal vitamin D deficiency alters immune cell proportions in offspring by changing the transcriptional properties of genes downstream of vitamin D receptor signaling in hematopoietic stem and progenitor cells of both the fetus and adults. Our results suggest the role of cellular differentiation properties of the hematopoiesis as the long-term memories of prenatal exposure at the adult stage. Moreover, further investigations of the associations between maternal vitamin D levels and cord blood immune cell profiles from 75 healthy pregnant women and their term babies also confirm that maternal vitamin D levels in the second trimester significantly affect immune cell proportions in the babies. This highlights the importance of providing vitamin D supplementation at specific stages of pregnancy.

## INTRODUCTION

Vitamin D, a micronutrient/hormone, regulates transcription by binding to its nuclear receptor, the vitamin D receptor (VDR). Humans can obtain vitamin D through photosynthesis on the skin and food intake. Approximately 50% to 90% of vitamin D is synthesized on the skin via sunlight exposure, while the remainder comes from the diet. Cutaneous vitamin D synthesis depends on environmental factors (geographic latitude, season, and amount of air pollution), skin type, clothing habitats, and lifestyle^1^. Despite country-level vitamin D fortification programs in many countries, the prevalence rate of vitamin D deficiency, especially in reproductive-age women^2–7^, is still high worldwide^8–14^. The most severe consequence of vitamin D deficiency is rickets or impaired bone formation (reviewed by Holick^15^). Besides its importance in bone formation, VDR signaling regulates gene transcription in nearly every tissue in our bodies, including the brain, heart, muscle, kidney, and immune system (reviewed in Pike et.al^16^). During development, the micronutrient status of offspring is entirely dependent on the status of the mother; therefore, developing embryos are vulnerable to adverse micronutrient conditions of deficient mothers^17–21^. To study the adverse consequences of prenatal vitamin D exposure, animal models with maternal dietary manipulations^22–35^ and knockout mouse model studies^18,36–42^ have been utilized. Studies using knockout mice models have shown that depletion of VDR causes a rickets-like phenotype after weaning. Both maternal dietary manipulation and knockout mouse models revealed that prenatal vitamin D deficiency can lead to immune defects in offspring. Epidemiological studies in humans have demonstrated that prenatal vitamin D deficiency has been associated with susceptibility to a number of immune-related diseases affecting their children, including asthma^43,44^, multiple sclerosis^45^, and type I diabetes^46–48^. These findings indicate that prenatal vitamin D deficiency disturbs immune cell development. In addition, it is possible that the hematopoietic system poses a long-term memory of exposure to vitamin D deficiency during development. However, this memory mechanism has not been extensively studied.

In this work, we show that prenatal vitamin D deficiency alters immune cell proportions of offspring during adulthood, attributed to cell fate decisions influenced by transcriptional alterations of VDR signaling pathway genes. Our data also supports that changing the cell fate of stem cells is the key component of the long-term effects of maternal vitamin D deficiency on the hematopoietic system of offspring.

## RESULTS

### Lack of impact of maternal vitamin D deficient diet feeding on offspring bone development

We summarized the dietary intervention strategy in **Fig. 1**. We randomly assigned six weeks old female C57BL/6J mice to a vitamin D-sufficient diet (VDsuf,1.0 IU/g vitamin D) and a vitamin D-deficient diet (VDdef, 0.0 IU/g vitamin D) for five weeks prior to mating with control diet-fed C57BL/6J male mice. The daily food intakes of each diet were comparable between the VDsuf and VDdef. To prevent hypocalcemia, the VDdef group received 1.5% calcium gluconate in their drinking water during the VDdef diet feeding. After five weeks of feeding VDdef, serum vitamin D (25-hydroxyvitamin D_3_, 25(OH)D_3_) concentrations of the VDdef-fed females reached the vitamin D deficient threshold (5.1±3.8 ng/mL, n=8), which is about 6.5 times lower than that of the VDsuf-fed females (33.0±5.44 ng/mL, n=10) (**Supplementary Fig. 1A**).

**Fig. 1:**
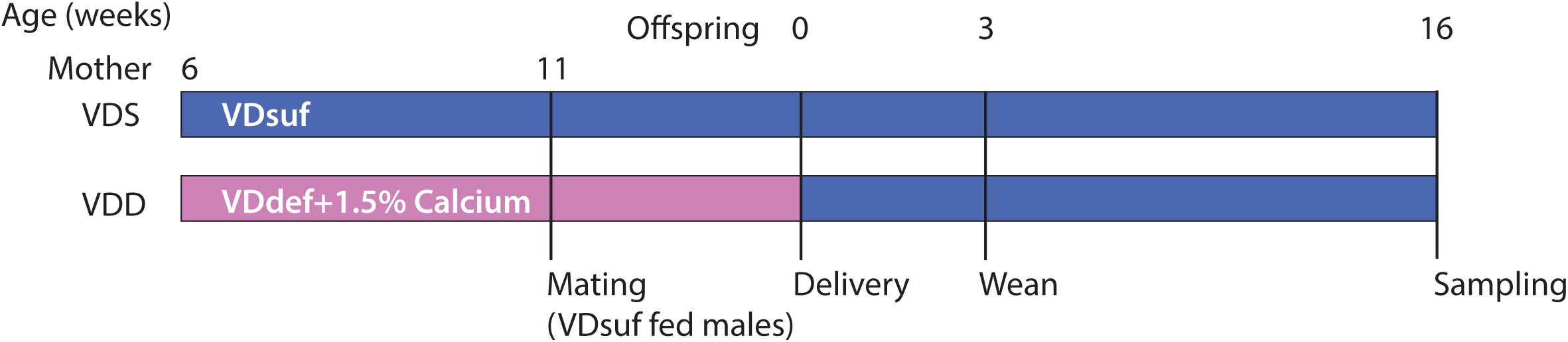
Study design. Six-week-old female C57BL/6J mice were randomly assigned to the VDsuf or VDdef diet and fed the assigned diet for five weeks before mating with a control diet-fed male. 1.5% calcium gluconate water was supplemented to VDdef diet-fed group. After delivery, all F0 mice were fed VDsuf diets. The offspring were fed VDsuf diet after weaning and maintained the diet until the sampling.

After delivery, all female mice were fed VDsuf. While the number of offspring per dam was smaller in VDdef-fed females (mean = 3.82, standard deviation (sd) = 2.8, n=16) than in VDsuf-fed females (mean=4.7, sd=1.7, n=10), the difference was not significant (p=0.32). We did not observe any obvious adverse effects of VDdef diet feeding on these female mice prior to or during their pregnancy. All offspring were fed VDsuf after weaning and maintained the diet until the sampling. The growth of VDdef-fed female offspring (VDD) and VDsuf-fed female offspring (VDS) based on their body weight were comparable for all of the time points we measured: postnatal day 1 (PD1) (n=9 VDS and n=11 VDD), 5, 9, and 13 weeks of age (n=40 VDD and n=27 VDS) (**Supplementary Fig. 1B and 1C**). While VDD males were slightly smaller than VDS males, as Seipelt et al. reported^49^, the difference was not statistically significant at all time points. This discrepancy could be attributed to the differences in diet used or calcium supplementation to VDdef-fed mothers. The serum vitamin D concentrations of VDD were lower than the detection limit (0.6 ng/ml, n=3 per group) at PD1 (**Supplementary Fig. 1A**). While the serum vitamin D concentration of VDD was still significantly lower than that of VDS (n=5 per group) at five weeks of age, their status was not deficient anymore (**Supplementary Fig. 1A**). We collected tissues and weighted liver, lung, kidney, thymus, spleen, and heart from the VDD and VDS at 16 weeks of age (16 wks) (n=4-6 for each group). The average body weight adjusted tissue weights were not significantly different between the groups, except the heart was heavier in male VDD compared to male VDS (p=0.041) (**Supplementary Fig. 1D**). We also assessed the bone and tissue mineral density of the humerus of the left arm of the offspring using an X-ray CT system at 16 wks (n=5-6 males for each group). Bone volume, bone mineral content, bone mineral density, tissue mineral content, tissue mineral density, and bone volume fraction were assessed, and no significant alterations were observed (**Supplementary Fig. 1E**).

### Prenatal vitamin D deficiency decreases CD4+ and CD8+ T cell proportions of the offspring in peripheral blood and spleen

We collected peripheral blood from VDD and VDS and compared the immune cell profiles at 16 wks (n=11 per group). We observed a significant reduction of CD4+ T cells (28.2% decrease at p=0.0018) and CD8+ T cells (14.7% decrease at p=0.037) in VDD males compared to VDS males (**Fig. 2a**). This reduction was not observed in females (**Supplementary Fig. 2**). We also compared the immune cell profiles of spleen (n=6 per group) and observed the significant reduction of CD4+ T cells (18.4% decrease at p=0.0035) and CD8+ T (11.1% decrease at p=0.026) cells in VDD (**Fig. 2b**). Representative flow cytometry traces are shown in **Supplementary Fig. 3**.

**Fig. 2:**
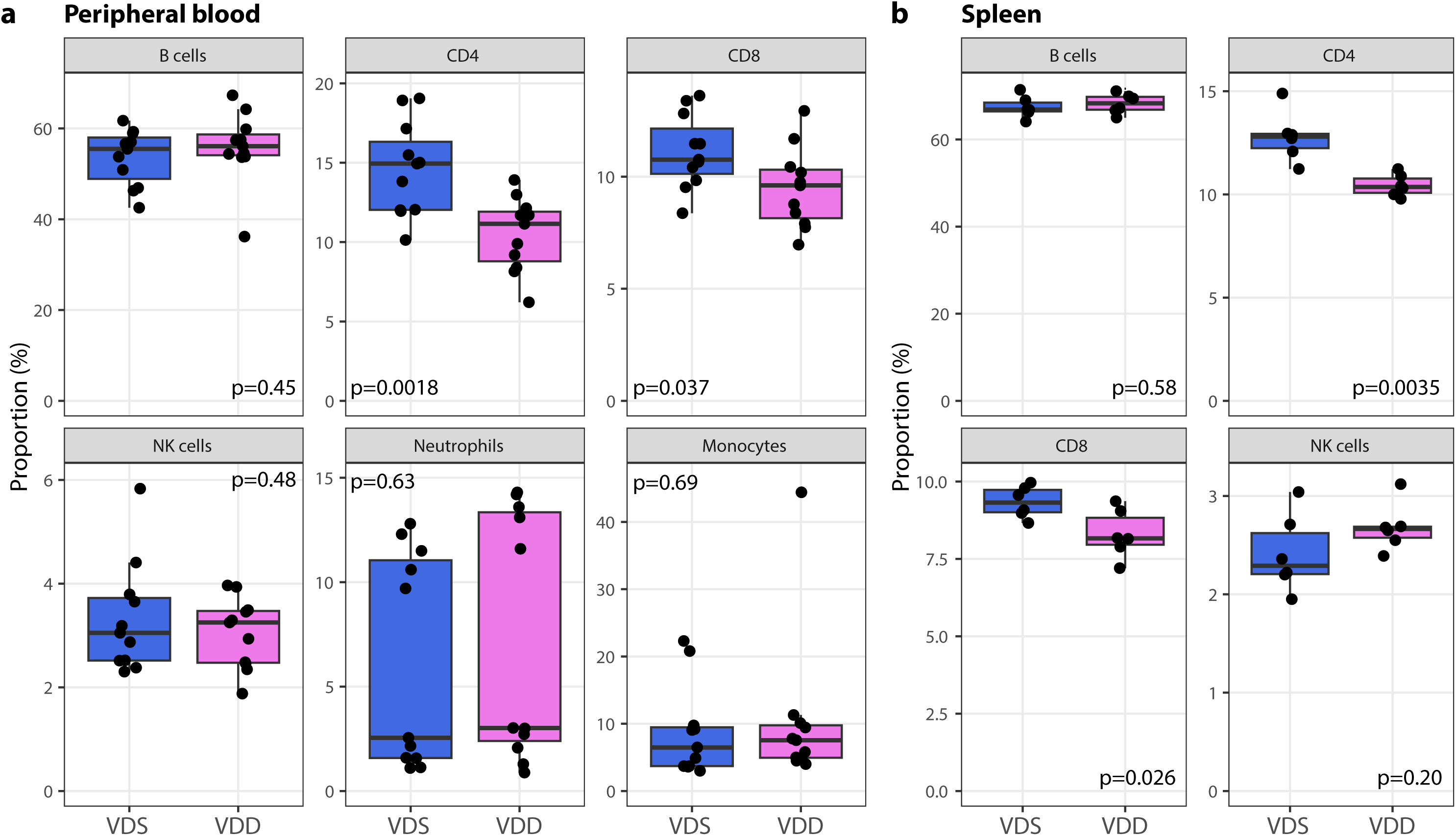
Prenatal vitamin D deficiency reduces CD4+ and CD8+ T cells in the periphery at the adult stage. T cell proportions in the peripheral blood (**a**) and spleen (**b**) were significantly decreased in VDD offspring. Each dot on the plot represents a sample (blood, n=11 per group; spleen, n=6 per group). The box shows the range between the first and third quartiles. The upper and lower whiskers represent 1.5 times the interquartile range, while the black bars indicate the median. The values shown in the plot are p-values calculated using Student’s t-test.

### The reduced proportion of lymphocytes in the periphery reflects the cellular composition changes in the bone marrow

During hematopoiesis, hematopoietic stem cells (HSCs) undergo differentiation into three multipotent progenitor cells (MPP2, MPP3, and MPP4) in the bone marrow in adults^50^. MPP2 and MPP3 are further differentiated primarily into the myeloid lineage, while MPP4 is into the lymphoid lineage. MPP2 is biased toward producing megakaryocytes and erythrocytes, and MPP3 towards granulocytes and macrophages^50^. To test if the T cell reductions in the periphery are reflected in the alterations in bone marrow, we collected mononucleated cells from bone marrow from the offspring at 16wks (n=9-10 per group). The total number of bone marrow mononucleated cells in VDD was significantly decreased compared to VDS (3,354,105±439,109 and 2,215,799±353,525 in VDS (n=10) and VDD (n=10), respectively, p=0.000013, **Fig. 3a**). We, then analyzed the proportions of HSCs, MPPs, and hematopoietic progenitor cells using the previously reported definitions^50,51^; lineage negative (Lin-, CD3-/CD4-/CD8-/B220-/Ter119-/CD11b-/Gr-1-/CD127-), Lin-/Sca-1+/c-Kit+ (LSK), long-term hematopoietic stem cell (LT-HSC, LSK/Flk2-/CD150+/CD48-), short-term hematopoietic stem cell (ST-HSC, LSK/Flk2-/CD150-/CD48-), multipotent progenitor 2 (MPP2, LSK/Flk2-/CD150+/CD48+), multipotent progenitor 3 (MPP3, LSK/Flk2-/CD150-/CD48+), multipotent progenitor 4 (MPP4, LSK/Flk2+), common myeloid progenitor cells (CMP, Lin-/c-Kit+/CD34+/CD16/32-), common lymphoid progenitor cells (CLP, Lin-/c-Kit+/Flk2+/CD127+), granulocyte-monocyte progenitor cells (GMP, Lin-/c-Kit+/CD34+/CD16/32+), megakaryocyte-erythrocyte progenitor cells (MEP, Lin-/c-Kit+/CD34-/CD16/32-), and Lin-/CD127+ (early T cell progenitor). We observed a significant reduction (48.9% decrease) in the absolute number of LSK (565,312±153,178 and 289,052±91,984 in VDS and VDD, respectively, p=0.00023, **Fig. 3b**). While the p-value (p=0.078) didn’t pass our significant threshold, the proportion (%) of LSK in the live bone marrow mononucleated cells was also decreased in VDD mice (0.19±0.04% and 0.16±0.05% in VDS (n=10) and VDD (n=9), respectively, **Supplementary Fig. 4a**). As the absolute number of LSK decreased in VDD, the absolute numbers of HSCs, MPPs, and progenitors were also decreased (**Fig. 3c and d**). Among the MPPs, MPP4 was the only MPP that significantly reduced in both absolute numbers (413,267± 127,529 and 190,922± 49,401 in VDS (n=10) and VDD (n=9), respectively, p=0.0027, **Fig. 3c**) and the proportion of the live bone marrow mononucleated cells (0.14±0.03% and 0.10±0.02% in VDS and VDD, respectively, p=0.0096, **Supplementary Fig. 4b**). Among the progenitor cells, the absolute numbers of CMP, GMP, CLP, and Lin-/CD127+ were significantly reduced in VDD (n=9) compared to VDS (n=9) (**Fig. 3d**). Interestingly, while the proportional averages of Lin-/CD127+ (p=0.07) and CLP (p=0.19) were decreased, the proportion of GMPs was increased in VDD compared to VDS (50.1% increase, p=0.001, **Supplementary Fig. 4c**). Representative flow cytometry traces are shown in **Supplementary Fig. 5**.

**Fig. 3:**
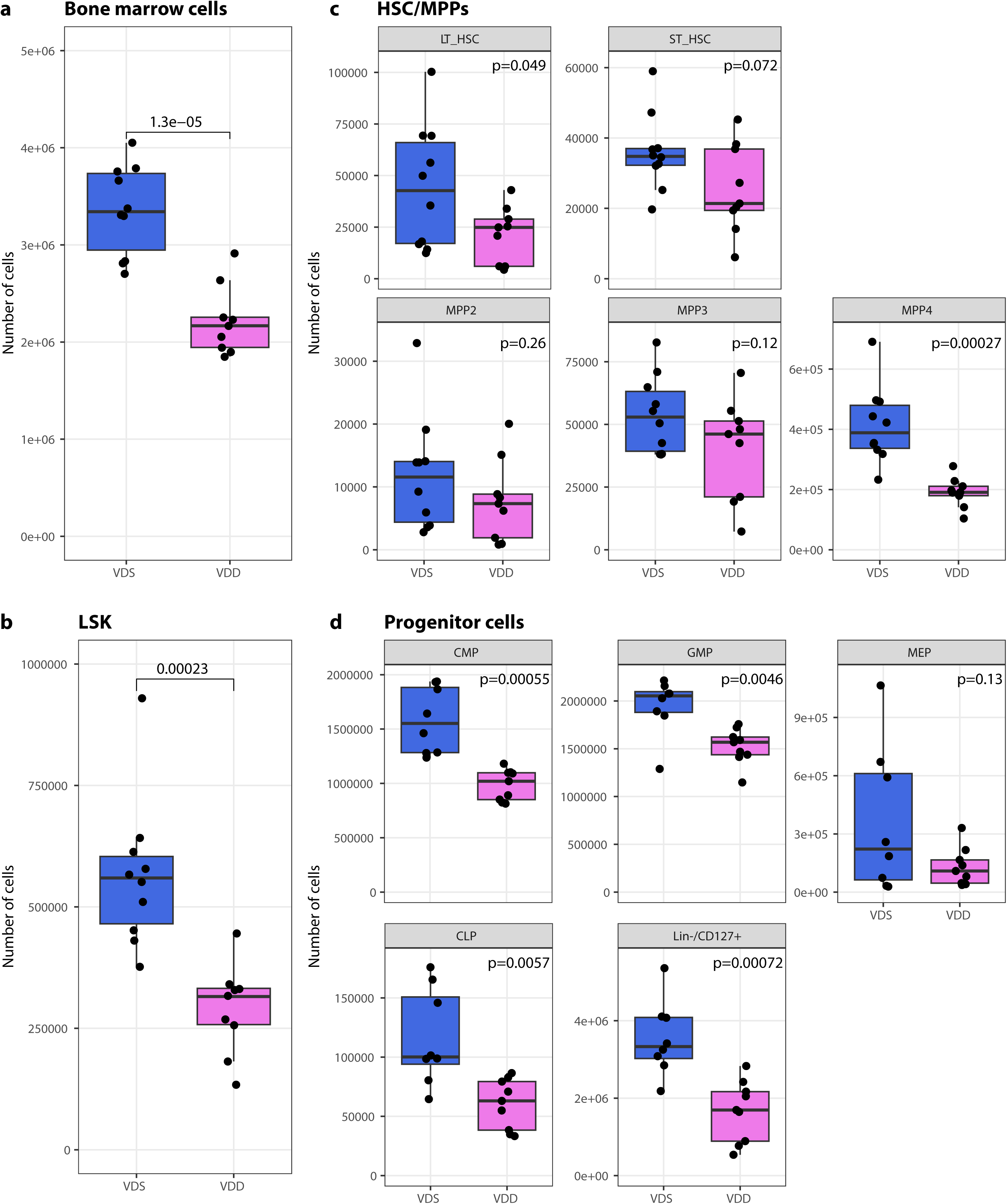
Prenatal vitamin D deficiency decreased the number of bone marrow cells and LSKs and concomitantly reduced lymphoid lineage cells at the adult stage. Prenatal vitamin D deficiency reduced the total number of bone marrow cells (n=9 for VDD and n=10 for VDS) (**a**) and the number of LSKs in bone marrow (n=9 per group) (**b**). The differences in total HSC, long-term and short-term HSC, and three MPPs in the bone marrow (n=9 for VDD and n=10 for VDS) (**c**) and hematopoietic progenitor cells in the bone marrow (n=9 per group) of VDD and VDS mice are displayed in the box plots (**d**). A significant decrease in MPP4, CLP, and Lin-/CD127+ cells was observed in VDD mice, indicating that prenatal vitamin D deficiency disproportionately affects the production of hematopoietic lineages. Each dot on the plot represents a sample. The box shows the range between the first and third quartiles. The upper and lower whiskers represent 1.5 times the interquartile range, while the black bars indicate the median. The values shown in the plot are p-values calculated using Student’s t-test.

### Prenatal vitamin D deficiency alters gene expression profiles of MPP4 cells

We performed transcriptome analysis on bone marrow MPP4 from VDD and VDS to see the transcriptional alterations at 16 weeks of age (n=3 per group). The sequencing status and quality of each sample are summarized in **Supplementary Data 1**. All samples have passed primary quality checks. A hierarchical clustering analysis showed a clear dissociation, suggesting genome-wide transcriptional alterations in VDD (**Fig. 4a**). We identified 612 differentially expressed genes (DEGs) with at least 1.2-fold change and a false discovery rate adjusted p-value (FDR-adj p-value) <0.05 (**Fig. 4b, Supplementary Data 2**). Of those, 345 were upregulated, and 267 were downregulated. The Gene Ontology (GO) enrichment analysis of upregulated genes revealed enrichment of genes related to leukocyte migration and chemotaxis (**Fig. 4c left**), while downregulated genes showed enrichment in the homeostasis of cell numbers and regulation of hemopoiesis (**Fig. 4d left, Supplementary Data 3**). The Gene-Concept Network analysis revealed that hematopoietic transcription factors, including *Lmo2*, *Tal1*, *Gata2*, and *Mpl*, were downregulated and were found to overlap in the top five enriched GO terms (**Fig. 4d right**). Transcription factor (TF) Perturbations Followed by Expression analysis (https://maayanlab.cloud/Enrichr/) revealed that the detected DEGs were enriched in the genes perturbed in LSK of the hematopoietic transcription factor knockout models in the same directions. The upregulated genes were enriched in NFIX (GSE45492) ^52^ and SRF (GSE23556) ^53^ knockouts upregulated genes (adjusted p-value=3.92E-47 and 9.15E-29, respectively) and downregulated genes were enriched in NFIX and SRF knockout downregulated genes (adjusted p-value=1.98E-36 and 4.55E-12, respectively) (**Supplementary Fig. 6**). Gene Set Enrichment Analysis (GSEA) showed leukocyte differentiation (GO:0002521, enrichment score = 0.385, q-value= 0.010), lymphocyte differentiation (GO:0030098, enrichment score = 0.412, q-value= 0.026), T cell differentiation (GO:0030217, enrichment score =0.435, q-value=0.023) were enriched (**Supplementary Data 4**). This result suggests that prenatal vitamin D deficiency induces long-term transcriptional alterations in hematopoietic stem cells that persist into adulthood.

**Fig. 4:**
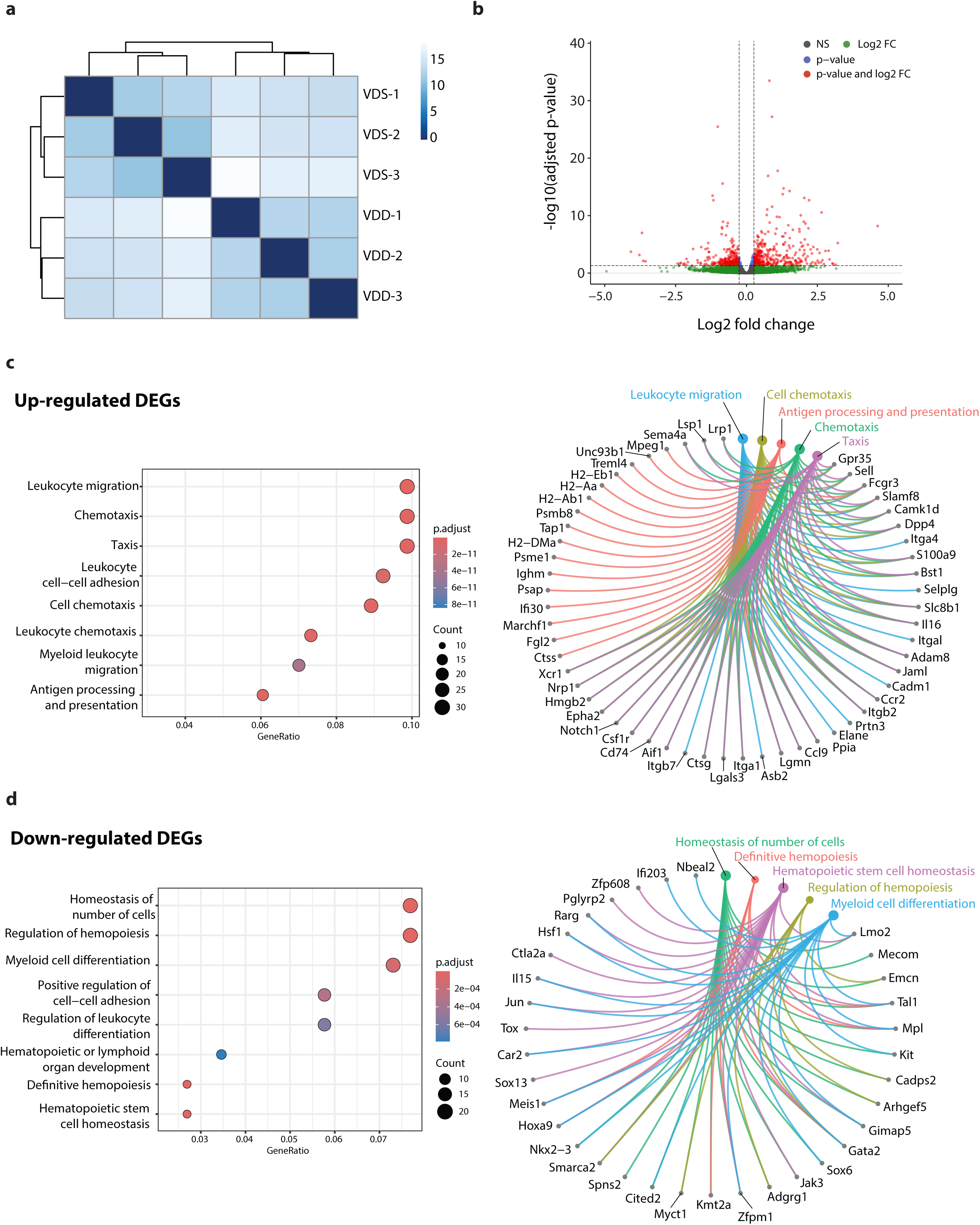
Bulk RNA-seq analyses reveal the transcriptional alterations in MPP4. Bulk RNA-seq analysis identified significant differences in the transcriptional profiles of MPP4 between VDD and VDS (n=3 per group). (a) The heatmap and dendrogram show a distinct clustering between VDD and VDS. (**b**) The volcano plot displays the −log10 adjusted p-values and log2 fold-change differences of the identified DEGs. Each dot represents a gene, and significant DEGs are indicated by red dots. (**c**) The GO enrichment analysis reveals that the up-regulated DEGs between VDD and VDS MPP4 are enriched in Leukocyte migration and chemotaxis-related genes. The dot plot shows the significance and the gene ratios of the top eight GO terms (left), and the Cnet plot indicates the identified DEGs of the top five GO terms (right). (**d**) The GO enrichment analysis reveals that the down-regulated DEGs between VDD and VDS MPP4 are enriched in the regulation of hematopoiesis-related genes. The dot plot shows the significance and the gene ratios of the top eight GO terms (left), and the Cnet plot indicates the identified DEGs of the top five GO terms (right).

### Prenatal vitamin D deficiency alters cellular compositions of the embryonic day 14.5 (e14.5) embryonic liver, suggesting immune cell proportion changes start during development

To assess if the cell composition alteration started at the embryonic stage, we performed single-cell RNA-seq (scRNA-seq) on the e14.5 fetal liver of both VDS and VDD embryos using the 10x Genomics Chromium platform (n=3 per group). Multiplexing with cell hash antibodies was used to reduce the technical batch effect. The obtained sequences were aligned by the 10x Genomics software, CellRanger, and the matrix was demultiplexed and analyzed by Seurat, an R-package ^54–56^. After eliminating low-quality cells (<1000 genes/cell, <5000 reads/cell, and >10% mitochondrial reads/cell) and cells without cell hashing information, we have identified 21 different cell clusters in a total of 6947 cells. We identified the cell types of each cell cluster based on the marker gene expression status (**Fig. 5a, Supplementary Data 5**). We detected three HSC/MPP populations, lineage-specific progenitor cells, erythroid lineage cells, and hepatoblast cells in e14.5 liver. Concordant with the previous report^57^, more than half of the cells were identified as erythroid lineage cells. The cellular composition analysis showed that one of the HSC/MPP populations (HSC/MPP 1) was significantly increased in VDD embryos, and two erythroid lineage cells (Early Erythroid 1 and Erythroid 3) were decreased (**Fig. 5b**), suggesting the cell composition changes started at least at the embryonic stage. A pseudo bulk RNA-seq analysis on the scRNAseq datasets showed that down-regulated genes were enriched in the genes regulated by hematopoietic system transcription factors (**Fig. 5c, Supplementary Data 6-8**). Among these transcription factors, *Tal1*, *Lmo2, Meis1*, and *Erg* were also identified as VDD downregulated genes in MPP4 at the adult stage (**Supplementary Data 2**). To test whether vitamin D alters the expression of these genes, we measured the expression of *Tal1*, *Lmo2,* and *Erg*, the known hematopoietic transcription factors, by quantitative RT-PCR on an embryonic stem cell-derived hematopoietic progenitor cell line, HPC-7^58^, after treating 1-alpha-25-dihydroxyvitamin D_3_, a ligand of VDR, or ethanol (solvent) for 24 hours by quantitative RT-PCR (**Fig. 4d**). We observed a significant increase of *Lmo2* (p=0.034) and *Erg* (p=0.005) expression. This result suggests that the gene expression alterations observed at e14.5 were attributed to the dysregulation of hematopoietic transcription factors, such as *Lmo2* and *Erg,* regulated by VDR signaling pathways.

**Fig. 5:**
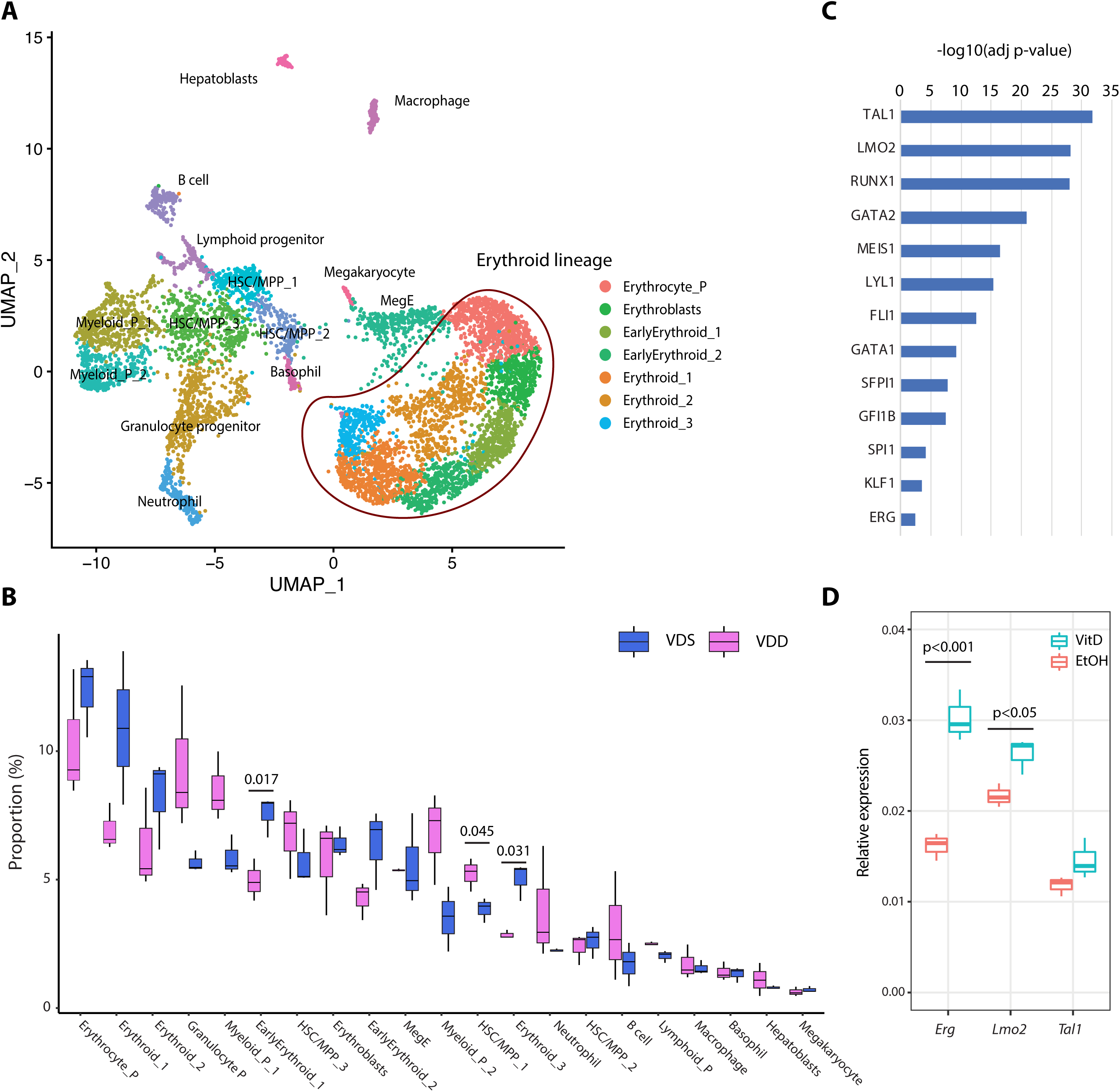
Prenatal vitamin D deficiency alters cellular compositions of the embryonic liver, suggesting immune cell proportion changes start during development. (**a**) UMAP representation of single-cell RNA-seq gene expression data and cellular lineage identification of E14.5 fetal liver (n=3 per group). (**b**) The boxplots indicate prenatal vitamin D deficiency alters cellular compositions of E14.5 fetal liver. (**c**) Genes downregulated in VDD E14.5 fetal liver are enriched in the genes regulated by hematopoietic transcription factors. (**d**) Treating HPC7 cells with 1-alpha-25-dihydroxyvitamin D_3_ significantly increases gene expression levels of *Erg* and *Lmo2*, suggesting these genes are regulated by VDR (n=3 per treatment group).

### Maternal serum vitamin D status in the second trimester is positively associated with the CD8+ T cell proportion in the cord blood

To test the associations between maternal vitamin D levels and immune cell proportions of offspring, we assessed 75 pregnant Japanese women who were recruited as participants of The Birth Cohort Gene and Environment Interaction Study of TMDU (BC-GENIST) project at the Tokyo Medical and Dental University, Bunkyo, Tokyo, Japan^59,60^. The study was approved by the Institutional Review Board of Tokyo Medical and Dental University (No. G2000-181, 29 July 2014). We measured the concentration of serum 25(OH)D_3_ at two-time points: time point 1 (T1), intended to represent the second trimester from week 10 to week 29, and time point 2 (T2), to represent the third trimester from week 33 to week 40 of gestational age. The immune cell proportions were estimated from the bulk DNA methylation profiles of the cord blood of the fetus using a Bioconductor package FlowSorted.CordBloodCombined.450k^61–64^. We excluded one participant who did not have T1 serum 25(OH)D_3_ status from the analysis. The demographic, clinical, and phenotypic information for the study participants is provided in **Table 1**. All participants are healthy pregnant Japanese women without smoking or drinking during their pregnancy. No participants had hypertension. The average maternal age at delivery was 34.2 (sd 4.0) years old, with pre-pregnancy BMI 17.1-29.2 (average 20.73, sd 2.6). 70.3% of women took prenatal vitamin supplements, and the average estimated daily dietary vitamin D intake was 5.1 (sd 4.4) µg/day^65,66^. 34 offspring were males (45.9%), and the average gestational week of the delivery was 39.1 (sd 1.2). The average concentration of serum 25(OH)D_3_ was 25.2 (sd 11.8) ng/ml in the second trimester (average gestation week 19.0 (sd 4.5), T1) and 28.0 ng/ml (sd 14.4) in the third trimester (average gestation week 35.9 (sd 0.9), T2). 30 participants were vitamin D deficient (serum 25(OH)D_3_ <20 ng/ml) at T1 (40.5%), and 23 were vitamin D deficient at T2 (31.8%). Of those, eight participants were vitamin D deficient at both T1 and T2. While we observed significant associations between the serum 25(OH)D_3_ levels and sampling season (Summer and Winter) (p=0.047 and p<0.001, T1 and T2, respectively), no significant associations were observed in maternal age, pre-pregnancy BMI, sex of fetuses, prenatal vitamin supplements usage and dietary vitamin D intake. The results of the univariate model fitting are shown in **Supplementary Data 9**. To analyze the associations between maternal serum 25(OH)D_3_ levels and immune cell profiles of the cord blood, we calculated principal components (PCs) of immune cell profiles and assessed the contributions of each covariate. Gestation weeks at delivery showed the most significant contributions to PC1 (p= 0.0063 and adjusted r^2^=0.087) and PC2 (p= 0.000027 and adjusted r^2^=0.208), followed by maternal serum 25(OH)D_3_ at T1 to PC1 (p= 0.021 and adjusted r^2^=0.059) and being born in summer (p= 0.049 and adjusted r^2^=0.04) (**Fig. 6a**). We then assessed the associations of covariates to each cell type. We observed that the gestational weeks at delivery were negatively associated with the proportions of CD4 T cells (estimate −0.027 and p=0.00006) and B cells (estimate - 1.12 and p=0.00002) and positively associated with proportions of granulocytes (estimate 0.028 and p=0.0014). Maternal serum 25(OH)D_3_ at T1 was positively associated with proportions of CD8 T cells (estimate 0.0007 and p=0.0216) and monocytes (estimate 0.0005 and p=0.0349) and negatively associated with proportions of granulocytes (estimate −0.0024 and p=0.0365) (**Supplementary Fig. 7**). These associations were still significant after adjusting for the sex of the fetus and gestational weeks at delivery, season of T1, and the gestational week at T1 (**Fig. 6b, Supplementary Data 10**). These findings indicate that maternal vitamin D levels, especially in the second trimester, affect the immune cell proportions in the babies at birth.

**Fig. 6:**
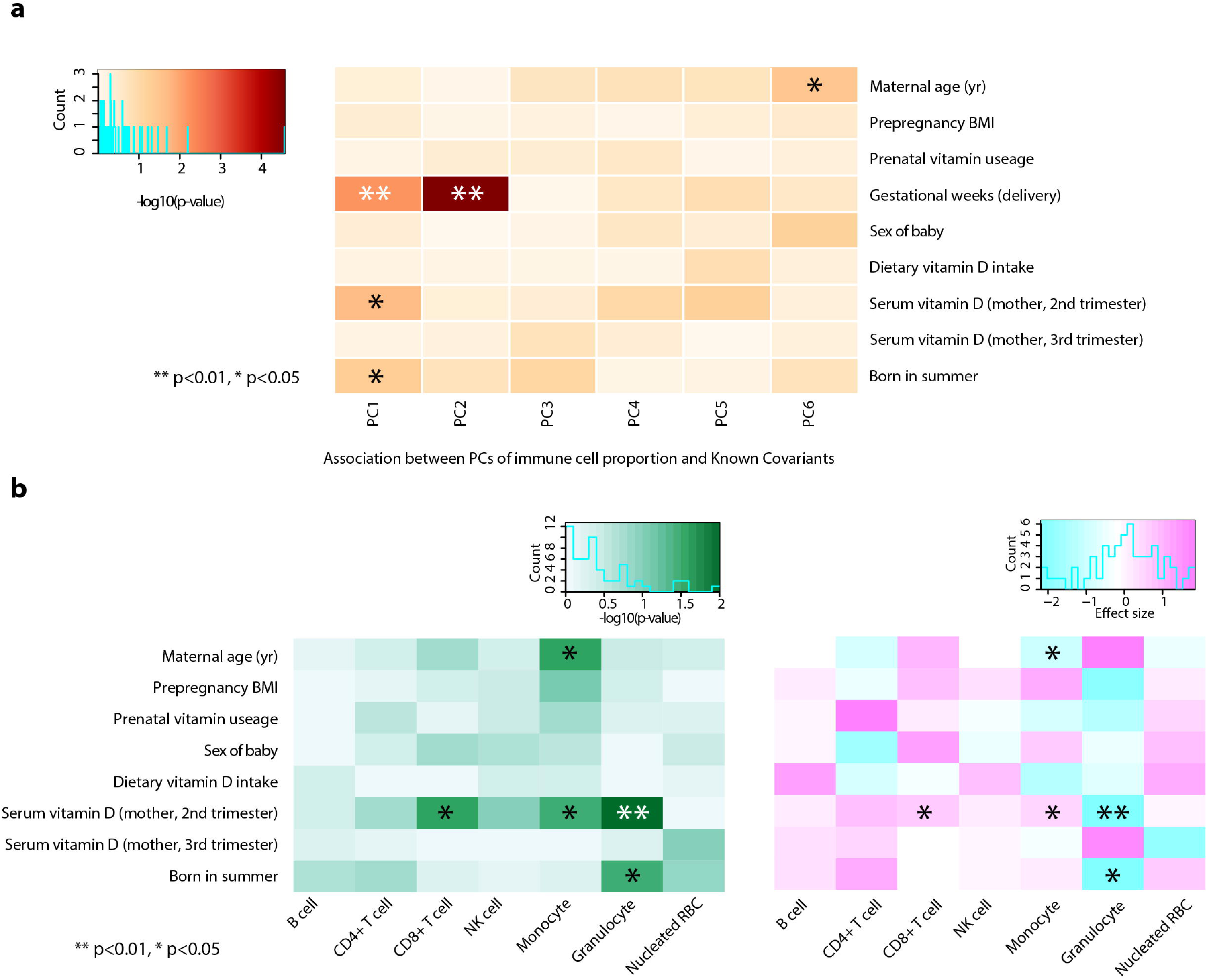
Maternal serum vitamin D status in the second trimester is positively associated with the CD8+ T cell proportion in the cord blood. (**a**) The heatmap shows that the gestational week at the delivery has the strongest associations with immune cell composition variations assessed by the principal component (PC), followed by maternal serum vitamin D (2^nd^ trimester) and being born in the summer season. (**b**) After adjusting for the sex of the fetus, gestational age, the season of T1, and the gestational week at T1, maternal serum vitamin D (2^nd^ trimester) maintains significant associations with immune cell composition, specifically positive association with proportions of CD8+ T cell and monocytes, and negative association with granulocytes. Asterisks indicate the significance (** p<0.01 and * p<0.05, Student’s t-test). The left panel shows −log10(p-value), and the right panel shows the direction of the associations.

**Table 1:** Demographic information of BC-GENIST cohort.

|  |  |  |  |
| --- | --- | --- | --- |
| Total N (%) |  |  | 74 (100.0) |
|  | Maternal age (yr) | Mean (SD) | 34.2 (4.0) |
|  | Pre-pregnancy BMI | Mean (SD) | 20.7 (2.6) |
|  | Prenatal vitamin usage, N (%) | Yes | 52 (70.3) |
|  | Fetal sex, N (%) | Female | 40 (54.1) |
|  | Gestational age at delivery (weeks) | Mean (SD) | 39.1 (1.2) |
|  | Gestational age at T1 | Mean (SD) | 19.0 (4.5) |
|  | Serum vitamin D concentration (ng/ml) at T1 | Mean (SD) | 25.2 (11.8) |
|  | Gestational age at T2 | Mean (SD) | 35.9 (0.9) |
|  | Serum vitamin D concentration (ng/ml) at T2 | Mean (SD) | 28.0 (14.4) |
|  | Estimated dietary vitamin D intake per day | Mean (SD) | 5.1 (4.4) |
|  | Maternal serum vitamin D deficiency levels, N(%) † | Deficient | 8 (10.8) |
|  |  | Insufficient | 20 (27.0) |
|  |  | Sufficient | 46 (62.2) |
| Estimated immune proportions from cord blood DNA methylation profiles |  |  |  |
|  | CD8 T cell | Mean (SD) | 5.23(3.35) |
|  | CD4 T cell | Mean (SD) | 19.25(7.15) |
|  | NK cell | Mean (SD) | 0.71(1.24) |
|  | B cell | Mean (SD) | 6.67(2.84) |
|  | Monocyte | Mean (SD) | 3.15(2.27) |
|  | Granulocyte | Mean (SD) | 61.46(9.2) |
|  | Nucleated red blood cell (nRBC) | Mean (SD) | 4.09(3.26) |

†: Deficient, both T1 and T2 serum vitamin D concentrations were less than 20 ng/ml; insufficient, one of the measurements was less than 20 ng/ml and the other was less than 30 ng/ml

## DISCUSSION

In this study, we found that prenatal vitamin D deficiency by maternal dietary intervention reduced peripheral T-cells in both the blood and spleen in adults and this reduction was linked to the decreased number of bone marrow cells and the proportional alterations of hematopoietic progenitor cells in the bone marrow. Our results demonstrate that cellular composition alterations could be the long-term memories of prenatal exposure at the adult stage.

Besides insufficient nutritional intake or photosynthesis on the skin, genetic mutations significantly contribute to vitamin D deficiency. In humans, mutations in genes of the vitamin D biosynthetic pathway are the most common cause of heritable vitamin D-dependent rickets (VDDR), which can be treated with active form vitamin D supplementation, whereas VDR mutation results in hereditary vitamin D-resistant rickets (HVDR), which could partially be ameliorated with Ca2+ supplementation^67,68^. Animal models with gene knockout have been studied to understand the functions of genes and diseases associated with them. A VDR Knockout mouse model study revealed that VDR null mice are developed without any significant defects before birth; however, they developed a rickets-like phenotype after weaning, and most of the animals died within 15 weeks after birth^40^, suggesting VDR is critical in the growth and bone formation in the post-weaning stage. Subsequent studies using the same VDR knockout mouse line feeding normal chow diet identified that while VDR knockout mice showed immune defects, no significant differences in the numbers or percentages of red and white cells compared to their wildtype littermates were observed^36,37^. Interestingly, correcting hypocalcemia by feeding a lactose-rich or polyunsaturated fat-rich diet was able to restore the immune abnormalities observed in VDR knockout mice^37^. Another research group independently developed another VDR knockout mouse line in C57BL/6 background and reported similar phenotypes, vitamin D-dependent rickets type II with alopecia after weaning^42^. Studies using this mouse line revealed that this line also developed immune abnormalities(39,69). Additionally, the nonobese diabetic (NOD) mouse line with a VDR knockout showed that although diabetes onset is not accelerated by VDR deletion, NOD VDR knockout mice develop rickets and have lower numbers of natural killer T-cells and CD4+CD25+ T-cells^41^.

These findings from independent knockout mouse studies indicate that VDR-depleted animals are phenotypically normal at birth but develop hypocalcemia within the first month of life and immune abnormalities. The active form 1-α-25-dihydroxyvitamin D [1,25(OH)_2_D] is synthesized from its precursor 25 hydroxyvitamin D [25(OH)D] via catalytic action of the 25(OH)D-1alpha-hydroxylase [1α(OH)ase] enzyme, encoded by Cyp27b1. Mice deficient in 1α(OH)ase developed hypocalcemia, skeletal abnormalities characteristics of rickets, female infertile, and a reduction in CD4 and CD8 peripheral T lymphocytes after weaning^18^. A zebrafish study showed that loss of Cyp27b1-mediated biosynthesis by gene knockdown significantly reduced *runx1* expression and hematopoietic stem and progenitor cell productions^67^. In this study, we found that prenatal vitamin D deficiency-exposed offspring at the adult stage did not show significant alterations in bone morphology, suggesting that vitamin D depletion during development does not affect embryonic bone development and bone morphology later in life. On the other hand, the immune phenotype of prenatal vitamin D deficiency-exposed offspring we observed was much more severe than those of VDR knockout models and similar to the Cyp27b1 deletion models. Since we observed this phenotype in offspring at 16 weeks of age, while VDR knockout mice typically do not survive beyond 15 weeks^40^, age might be another factor in developing this phenotype. However, our findings suggest that the immune phenotypes in offspring exposed to prenatal vitamin D deficiency may be caused by a lack of ligands rather than receptor malfunctions during the developmental phase.

A large biological difference between our study and the Cyp27b1 deletion models is the availability of 1,25(OH)2D after weaning. All offspring were fed VDS diets after weaning, and the serum vitamin D status of the VDD offspring became normal at five weeks of age. Therefore, the alterations we observed at 16 weeks of age were not associated with the vitamin D deficiency status at the time of measurement. However, it does indicate that prenatal vitamin D deficiency irreversibly alters bone marrow development in the offspring. This aligns with a previous study conducted on rats^69^. The study found that the rats with prenatal vitamin D deficiency by maternal dietary manipulation showed significant reductions in the total number of nucleated bone marrow cells and in the colony-forming unit (CFU) content with a corresponding increase in cell cycle rate^69^. The authors measured the CFU content by transplanting nucleated bone marrow cells to AJ mice purchased from Jackson Laboratory (vitamin D-sufficient diet-fed mice) and found that while 1,25(OH)2D3- or 24,25(OH)D3-treatment on vitamin D deficient rats in postnatal life corrected the serum calcium and phosphate, the treatments did not reverse the CFU content alterations. Their findings clearly demonstrate that prenatal and early-life vitamin D deficiency could cause irreversible effects on HSC development. However, we cannot fully rule out the possibility that alterations in hematopoietic stem cell niches might also be irreversibly altered. HSC niches in different developmental stages play critical roles in the maintenance and differentiation of HSCs (reviewed by Gao et al.^70^). It would be beneficial to explore this possibility further in upcoming research.

As a transcriptional regulation mechanism, 1,25(OH)2D binds to the VDR, which forms a heterodimer with the retinoid X receptor and regulates gene expression by binding to vitamin D responsive elements in genomic DNA. In this study, the transcription factor network analysis on e14.5 fetal liver scRNA-seq revealed that *Tal1*, *Lmo2*, *Runx1*, *Gata2, and Erg* downstream genes were significantly enriched in VDD downregulated genes. We confirmed that the expressions of *Erg* and *Lmo2* are regulated by VDR, as the treatment with 1-alpha-25-dihydroxyvitamin D3 (1,25(OH)_2_D3) led to an increase in gene expression in fetal hematopoietic progenitor cells, HPC7. A mutant and chimeric embryo study reported that ERG is a direct upstream regulator of *Runx1* and *Gata2* in fetal livers^71^, suggesting prenatal vitamin D deficiency exposure perturbs the transcription profiles of the hematopoietic cells during development via modulating hematopoiesis transcription factors regulated by VDR. Interestingly, this transcription factor expression perturbation persists in the adult stage even though these mice were fed a VDsuf diet and their vitamin D was fully recovered. Using bulk RNA-seq analysis, we identified 612 DEGs between VDD and VDS MPP4. Downregulated DEGs were enriched in hematopoietic-related pathways, and many hematopoietic transcription factors, including *Tal1*, *Lmo2*, *Runx1*, *Gata1*, and Erg, were downregulated. Gene Enrichment Analysis showed that the identified DEGs were enriched in the DEGs of *Nfix* and *Srf* knockout mouse LSKs in the same direction. *Nfix* and *Srf* are hematopoietic transcription factors. *Nfix* is associated with hematopoietic stem and progenitor cells (HSPCs) survival, and *Srf* regulates hematopoietic stem cell adhesions. Although both *Nfix* and *Srf* were not identified as DEGs, their expression levels were lower in VDD compared to VDS. It has been reported that loss of *Nfix* expression in HSPCs of murine adult bone marrow is concomitant with the reduced expression of genes associated with HSPCs survival, such as *Erg*, *Mecom*, and *Mpl* ^52^. These same genes were also significantly downregulated in VDD MPP4. However, the authors reported no selective loss of B, T, or myeloid cells was observed in *Nfix*-depleted HSPCs ^52^. On the other hand, Ragu *et al.* reported that while depleting *Srf,* increased % of the LSK in the bone marrow, *Srf* knockout mice showed a significant decline in white blood cell counts, which was primarily due to decreased numbers of circulating B and T cells ^53^. Our results suggest prenatal vitamin D deficiency modulates the *Nfix* and *Srf* pathways in the long term.

Our study also revealed that higher maternal vitamin D levels during the second trimester were associated with increased proportions of CD8+ T cells and decreased granulocytes in cord blood in humans. The associations were maintained after adjusting for the season of the second trimester, the gestational week at the maternal blood draw, the sex of the fetus, and the gestational week at birth. Hematopoietic development consists of primitive and definitive hematopoiesis^72^. In mice, the definitive hematopoiesis starts in the aorta-gonad mesonephros (AGM) region at approximately E10.5, then moves to the liver around E12.5 and the bone marrow around E17.5 ^73,74^. After birth, the bone marrow is the only site where HSCs are maintained and expanded. In human development, the transition of the hematopoietic stem cell production site from the fetal liver to bone marrow occurs in the second trimester^74,75^ when we observed significant associations. Recently, Elgormus et al. reported that newborn serum vitamin D levels were negatively correlated with neutrophil-to-lymphocyte ratios (NLR) in newborn babies^76^. The most common white blood cell is the granulocyte, composed of three distinct types: neutrophils, eosinophils, and basophils. Among them, neutrophil is the most abundant type. Our results and their finding concordantly indicate that the immune cell compositions, especially granulocyte (neutrophil) and lymphocyte, of the babies are influenced by vitamin D levels in early life. Of note, cord blood NLR has been proposed as an indicator for the diagnosis of early neonatal sepsis combined with other laboratory tests and clinical manifestations^77–79^. The associations between neonatal sepsis and cord blood vitamin D levels have been well reported. A meta-analysis of 18 studies revealed that low maternal and cord blood vitamin D levels were significantly associated with the incidence of neonatal sepsis^80^. Additionally, a study on 4,340 neonates appropriate for gestational age found a negative correlation between cord blood NLR and fetal malnutrition. This indicates that cord blood NLR could be utilized as a marker for fetal malnutrition^81^. Supplementing with Vitamin D during pregnancy may lower the risks of neonatal sepsis and other adverse outcomes.

In conclusion, this study demonstrates that prenatal vitamin D deficiency reduces the number and the proportion of MPP4 cells in the bone marrow, changing the expression status of the genes that may be directly and indirectly regulated by VDR, and this alteration results in reduced T cell proportions in peripheral blood and spleens of the offspring. The association between T cell proportion and maternal vitamin D status was also observed in our BC-GENIST mother-baby cohorts. We and others have reported that micronutrient deficiency changed the cellular compositions of adult mature organs and may contribute to the phenotypes using animal dietary manipulation models^21,35,82^. These findings indicate that the cell fate decision alteration of stem cells could be the key component of the long-term memory of prenatal micronutrient deficiency, which links to disease risks later in life. Nevertheless, this study has several limitations. A study conducted in Greece on mother-infant pairs revealed that insufficient levels of vitamin D (<50 nmol/L) in pregnant mothers were linked to an increased occurrence of micronucleated cells in binucleated T-cells^83^. Micronuclei are early indicators of genetic effects used to test the relationship between exposure to genotoxic substances and cancer. However, our study did not explore whether maternal VDD increases DNA damage risks in offspring. Sexual dimorphism is another covariate we must explore. Since female mice did not show significant alterations in immune cell proportions in adults, we only performed transcriptional analysis on male mice. Both males and females may have prenatal and neonatal immune cell proportional alterations. Furthermore, there could be transcriptional changes in certain genes in adult females, but these changes may not affect cell fate decisions. These possibilities should be further investigated in future studies.

## METHODS

### Maternal vitamin D deficiency mouse model

We purchased five-week-old C57BL/6J female mice from the Jackson Laboratory. After one week of acclimation, we started dietary manipulations at six weeks old. Female mice (F0) were fed vitamin D deficient (VDdef) or sufficient (VDsuf) diets for five weeks before mating with the control diet-fed male mice and throughout the subsequent pregnancy. To avoid the paternal VDdef treatment effects, the males did not stay in the female cage for more than three days. VDdef (0.0 IU/g vitamin D) and nutrient-matched VDsuf (1.0 IU/g vitamin D) diets were obtained from Research Diets Inc. (10 kcal%fat, 20 kcal%Protein and 70 kcal%Carbonate). The detailed ingredients of each diet are listed in **Supplementary Data 11**. VDdef-fed females were supplemented with 1.5% calcium gluconate water for drinking water. After delivery, all F0 mice were fed VDsuf diets. The offspring were fed VDsuf diet after weaning and maintained the diet until the sampling. All animal studies were approved by the Institutional Animal Care and Use Committee at the Albert Einstein College of Medicine (protocol # 20160710).

### Serum vitamin D level of mouse serum samples

Serum vitamin D (25(OH)D) levels of mouse serum samples were assessed by commercially available ELISA kits (Eagle Biosciences, Inc. Amherst NH, or Abcam, Cambridge, MA) according to the manufacturer’s instructions. The signal was detected with BioTek Synergy 4 Microplate reader (Agilent Technologies), and the results were analyzed using a 4-parameter logistic regression algorithm (http://www.elisaanalysis.com/app). The measurements were performed as duplicates.

### Immune cell profiling on peripheral blood and spleen

Immune cell profiling analysis was performed using flow cytometry (FACS Aria2, BD Biosciences) after fluorescent dye conjugate antibodies staining. The obtained data were analyzed with FlowJo_10.6.1_CL (https://www.flowjo.com/). Peripheral blood samples were collected using submandibular vein bleeding methods. The spleen was obtained after euthanizing the animals with carbon dioxide inhalation. The spleen samples were dissociated on 70 µm filters. The obtained single-cell suspensions were stained with antibodies of immune cell surface marker proteins to identify cell types. Representative flow cytometry traces are shown in **Supplementary Fig. 3**. The antibodies used in this study are listed in **Supplementary Data 12**.

### Isolating multipotent progenitor cells from bone marrow

Bone marrow cells were collected by crushing bones (tibia, Femur, iliac, sternum, and vertebrae). The red cells were lysed with lysis buffer (150 mM NH_4_Cl, 1 mM KHCO_3_, and 0.1 mM EDTA). HSC/MPP fractions were defined by the previously reported definition^50^. MPP4, lymphoid-primed hematopoietic multipotent progenitor cells were isolated for gene expression analysis. Representative flow cytometry traces are shown in **Supplementary Fig. 5**.

### Cell culture

Hematopoietic progenitor cell line HPC-7 was kindly gifted by Dr. Britta Will at Albert Einstein College of Medicine. HPC-7 cells were maintained at density of 1-10 x10^5^/ml in Iscove’s modified Dulbecco’s medium (Invitrogen) supplemented with 50-100 ng/ml of mouse stem cell factor (Gemini Bio-Products), 1 mM Sodium Pyruvate, 6.9 ng/mL α-Monothioglycerol (SIGMA-Aldrich), 5% of bovine calf serum and Penicillin-Streptomycin. To examine the effect of vitamin D treatment on gene expression levels, the HPC-7 cells were treated with 0.1 µM of 1alpha,25-Dihydroxyvitamin D3 (SIGMA-Aldrich) or 0.1% (vol/vol) ethanol (solvent) for 24 hours.

### RNA-seq library construction and sequencing

We performed RNA-seq on FACS isolate MPP4 cells (lymphoid primed-multipotent progenitor cells). Total RNA was isolated with AllPrep DNA/RNA micro kit (QIAGEN). After we depleted ribosomal RNAs from total RNA, we generated the RNA-seq libraries using KAPA RNA HyperPrep with RiboErase kit (Roche). The generated libraries were sequenced on an Illumina NOVA-seq sequencer (Novogene Co., Ltd., USA).

### RNA-seq alignment

After checking the quality of the sequencing files using FastQC^84^ and trimming low-quality reads and adapter sequences using Cutadapt^85^, the obtained sequences were aligned to the mouse mm10 reference genome with the gencode M15 gene annotation using STAR aligner^86^. The quality of the library was assessed with RSeQC^87^. The obtained transcript counts were analyzed with DESeq2. We identified significant differentially expressed genes (DEGs) that showed two times higher or lower expression with a false discovery rate-adjusted p-value less than 0.05. The detailed sequencing and alignment status are listed in **Supplementary Data 1**.

### Enrichment analysis for Gene Ontology (GO)

We used the Gene Ontology (GO) enrichment analyses and Gene Set Enrichment Analyses (GSEA) of a Bioconductor package ClusterProfiler^88^ to see the functional enrichment of DEGs. Q-values <0.05 were considered significant.

### Quantitative RT-PCR

Total RNA samples were isolated with AllPrep DNA/RNA micro kit, and cDNA libraries were synthesized using SuperScript III transcriptase (Invitrogen) with random hexamer. The real-time PCR was performed with Roche LightCycler 480 SYBR GREEN I Master mix on a LightCycler 480 system (Roche). Relative gene expression abundance between samples was calculated using the CT method and *Gapdh* as an internal control. The sequences of primers used in this study are listed in **Supplementary Data 13**.

### Single-cell RNA sequencing library preparation

We used e14.5 male mouse liver (n=3 per group) to study transcriptional alteration at single-cell resolution. The e14.5 mouse liver samples were dissociated on 70 µm filters, then each sample was stained with unique cell hashing antibodies (BioLegend). 3,000-4,000 cells per sample were targeted on the 10x Genomics Chromium platform. Single-cell mRNA libraries were built using the Chromium Next GEM Single Cell 3’ Library Construction V3 Kit, libraries sequenced on an Illumina NOVA-seq. Sequencing data were aligned to mm10 mouse reference using the Cell Ranger 3.0.2 pipeline (10x Genomics). Counting cell hashing tag were performed using CITE-seq Count version 1.3.4^89^.

### scRNA-seq data processing, batch correction, clustering, cell-type labeling, and data visualization

All scRNA-seq analysis and data visualization were performed using an R package, Seurat^54–56^. After demultiplexing based on the cell hashing tag information, low-quality cells (<1000 genes/cell, <5000 reads/cell, and >10% mitochondrial reads/cell) were eliminated from the further analyses. Data integration and identifying cell clusters were carried out after performing SCTransform^56^. Cell types of each cell cluster were identified based on the expression of the marker genes^90^. Proportions of assigned cell types were analyzed using Student’s *t*-test. P-values <0.05 were considered significant. Differentially expressed genes of each cell type between VDD and VDS were identified as at least 25% of cells expressed the gene, the log fold change greater than 0.5, and the FDR adjusted p-value less than 0.05. GO enrichment analysis was performed using a Bioconductor package ClusterProfiler^88^ with q-values <0.05 considered significant. The enrichment status of the transcription factors was assessed using the CHEA transcription factor targets dataset^91^ and TF Perturbations Followed by Expression functions provided by Enrichr (http://amp.pharm.mssm.edu/Enrichr)^92^.

### Measurement of bone mineral density

Bone mineral density was measured humerus of the offspring at dissection. After the dissection, the left arms (n=5-6 animals for each group) were scanned using an X-ray CT system, Inveon Multimodality scanner (Siemens). The CT x-rays were generated by an 80kV peak voltage difference between the cathode and tungsten target at 0.5 mA current and 200 millisecond exposure time. The arm samples were placed on the 38mm width bed tandemly. The CT field of view was 5.5 cm by 8.5 cm with an overall resolution without magnification of 60 microns. A Scout View was performed before the start of the CT Acquisition to ensure the correct positioning of the subject in the field of view. Image analysis was performed using MicroView (https://microview.parallax-innovations.com/).

### Serum vitamin D level of serum samples from pregnant women

Healthy pregnant women were recruited as participants in The Birth Cohort Gene and Environment Interaction Study of TMDU (BC-GENIST) project at the Tokyo Medical and Dental University, Bunkyo, Tokyo, Japan^59,60^. Written informed consent was obtained from the participants, and the study was approved by the Institutional Review Board of Tokyo Medical and Dental University (No. G2000-181, 29 July 2014). In this study, all participants were healthy Japanese females aged 27-42 years old without smoking or drinking alcohol during their pregnancy. We used the following demographic information of the participants in this study; maternal age at delivery, prepregnant body mass index, prenatal vitamin supplements usage, sex of the fetus, gestation weeks, and estimated daily vitamin D intake. The daily vitamin D intake was estimated 3-day food record questionnaire. Maternal serum samples were collected twice, around 20 and 36 gestational weeks, and aliquoted samples were stored at −150 °C until use. Serum 25(OH)D_3_ levels were measured using a modified LC-APCI-MS/MS method^93^. As previously described, this method involves the use of deuterated 25(OH)D_3_ (*d*_6_-25(OH)D_3_) as an internal standard compound and the selection of a precursor and product ion with an MS/MS multiple reaction monitoring (MRM) method. The internal standard *d*_6_-25(OH)D_3_ (0.5 ng/10 μL) was added to serum (40 μL) and precipitated with acetonitrile (200 μL). After evaporation of the supernatant, the precipitant was dissolved with ethyl acetate (400 μL) and distilled water (200 μL) with vigorous shaking. The ethyl acetate phase was removed and evaporated. Extracted vitamin D metabolites from serum were derivatized by 4-[2-(6,7-dimethoxy-4-methyl-3-oxo-3,4-dihydroquinoxalyl)ethyl]-1,2,4-triazoline-3,5-dione (DMEQ-TAD) to obtain high sensitivity by increasing ionization efficiency^94^. Separation was carried out using a reverse-phase C_18_ analytical column (CAPCELL PAK C_18_ UG120, 5 μm; (4.6 I.D. × 250 mm) (SHISEIDO, Tokyo, Japan) with a solvent system consisting of (A) acetonitrile, (B) distilled water (0–5 min A = 30%, 5–34 min (A) = 30 → 70%, and 34–37 min (A) = 70 → 100%) as the mobile phase and a flow rate of 1.0 mL/min. All MS data were collected in the positive ion mode, and quantitative analysis was carried out using MS/MS-MRM of the precursor/product ion for DMED-TAQ-25(OH)D_3_ (m/z; 746.5/468.1) and DMED-TAQ-*d*_6_-25(OH)D_3_ (m/z; 752.5/468.1) with a dwell time of 200 ms (AB Sciex LLC., Framingham, MA, USA).

### Estimation of immune cell profiles of human cord blood

We used a modified deconvolution approach to estimate the immune cell profiles of human cord blood from the bulk DNA methylation profiles^95^. Bulk DNA methylation profiles of cord blood were accessed using Infinium HumanMethylation450 BeadChip, and the estimation was performed using a Bioconductor package FlowSorted.CordBloodCombined.450k^61^.

### Statistical analysis

Mouse phenotype results were analyzed using Student’s *t*-test. The associations between maternal serum vitamin D levels and immune cell profiles in human cord blood were tested using one-way ANOVA. P-values <0.05 were considered significant. Differences in gene expression between VDD-F1 and VDS-F1 in RNA-seq were analyzed using DESeq2 with FDR-adjusted p-value <0.05. R v4.0 (https://www.r-project.org/) was used for most of the analyses.

## DESCRIPTION OF SUPPLEMENTARY INFORMATION

**Supplementary Fig. 1**: The effects of vitamin D deficient diet feeding on the mothers and the impacts of prenatal VDD on the growth and bone density of offspring at the adult stage.

**Supplementary Fig. 2**: Prenatal VDD effects on female offspring at the adult stage.

**Supplementary Fig. 3**: Gating and analytical strategies to assess the immune cell profiles in peripheral blood and spleen.

**Supplementary Fig. 4**: Prenatal vitamin D deficiency alters hematopoietic cell cellular compositions of bone marrow at the adult stage.

**Supplementary Fig. 5**: Gating and analytical strategies to assess hematopoietic stem cells, multipotent progenitor cells, and progenitor cells from bone marrow.

**Supplementary Fig. 6**: Differentially expressed genes (DEGs) in VDD MPP4 are enriched in DEGs of hematopoietic transcription factor knockout models.

**Supplementary Fig. 7**: The heatmaps indicate the association between known covariates and immune cell proportions.

## DESCRIPTION OF ADDITIONAL SUPPLEMENTARY INFORMATION

**Supplementary Data 1:** RNA-seq sequencing stats (VDD and VDS MPP4)

**Supplementary Data 2:** A list of differentially expressed genes between VDD MPP4 and VDS MPP4

**Supplementary Data 3:** The results of GO enrichment analysis on MPP4 GEGs

**Supplementary Data 4:** The results of Gene Set Enrichment Analysis on MPP4 DEGs

**Supplementary Data 5:** A list of the marker genes of each cell cluster found in E14.5 fetal liver scRNAseq analysis

**Supplementary Data 6:** A list of DEGs of each cell cluster found in E14.5 fetal liver scRNAseq analysis

**Supplementary Data 7:** A list of DEGs in pseudo bulk RNA-seq analysis of E14.5 fetal liver scRNAseq

**Supplementary Data 8:** The results of Enrichr analysis on differentially expressed genes identified in pseudo bulk RNA-seq analysis of E14.5 fetal liver scRNAseq

**Supplementary Data 9:** Summary of the associations of known covariates to the maternal serum vitamin D levels

**Supplementary Data 10:** A list of the significance of the contribution to the proportions

**Supplementary Data 11:** The detailed ingredients of each diet

**Supplementary Data 12:** A list of antibodies used in this study

**Supplementary Data 13:** A list of primer sequences used in this study

## DATA AVAILABILITY

The authors declare that all data supporting the findings of this study are available within the article and its supplementary information files, except for human cord blood analyses, which may contain sensitive information. All sequencing data, RNA-seq and scRNA-seq of this study, is deposited in NCBI’s Gene Expression Omnibus GEO database under the accession number GSE242043, and processed data and code used in this study are available upon request.

## ACKNOWLEDGMENTS

The authors thank Dr. Britta Will at Albert Einstein College of Medicine for generously providing the HPC7 cells. Additionally, we would like to acknowledge the MicroPET Facility, supported by The M. Donald Blaufox Laboratory for Molecular Imaging and NIH (1S10RR029545” MicroPET/SPECT/CT Animal Imaging Device”), the Flow Cytometry Core Facility, and the Genomic Core Facility at Albert Einstein College of Medicine.

## AUTHOR INFORMATION

### Authors and Affiliations

These authors contributed equally: Koki Ueda, Shu Shien Chin, Noriko Sato

**Department of Cell Biology, Albert Einstein College of Medicine, Bronx, NY, USA**

Koki Ueda, Ulrich G. Steidl

**Department of Blood Transfusion and Transplantation Immunology, Fukushima Medical University, Fukushima, Fukushima, Japan**

Koki Ueda

**Department of Food and Nutrition, Faculty of Human Sciences and Design, Japan Women’s University, Bunkyo-ku, Tokyo, Japan**

Noriko Sato

**Department of Microbiology & Immunology, Albert Einstein College of Medicine, Bronx, NY, USA**

Shu Shien Chin, Laurent Chorro, Gregoire Lauvau

**Graduate School of Medical and Dental Sciences, Medical and Dental Sciences, Systemic Organ Regulation, Comprehensive Reproductive Medicine Tokyo Medical and Dental University, Bunkyo-ku, Tokyo**

Naoyuki Miyasaka

**Department of Genetics, Albert Einstein College of Medicine, Bronx, NY, USA**

Betelehem Solomon Bera, David Reynolds, Reanna Doña-Termine, John M. Greally, Masako Suzuki

**Department of Biotechnology, Faculty of Engineering, Toyama Prefectural University, Toyama, Japan**

Miyu Nishikawa

**Department of Pharmaceutical Engineering, Faculty of Engineering, Toyama Prefectural University, Toyama, Japan**

Kaori Yasuda

**Department of Radiology, Albert Einstein College of Medicine, Bronx, NY, USA**

Wade R Koba

**Ruth L. and David S. Gottesman Institute for Stem Cell Research and Regenerative Medicine, Albert Einstein College of Medicine, Bronx, NY 10461, USA**

Ulrich G. Steidl

**Department of Oncology, Albert Einstein College of Medicine – Montefiore Medical Center, Bronx, NY 10461, USA**

Ulrich G. Steidl

**Department of Pediatrics, Albert Einstein College of Medicine – Montefiore Medical Center, Bronx, NY 10461, USA**

John M. Greally

**Montefiore-Einstein Cancer Center, Albert Einstein College of Medicine – Montefiore Medical Center, Bronx, NY 10461, USA**

Ulrich G. Steidl

**Department of Nutrition, Texas A&M University, College Station, TX, USA**

Masako Suzuki

## Contributions

Conceptualization and methodology, G.L., U.G.S, J.M.G., and M.S.; investigation, K.U., N.S., S.S.C., M.N., K.Y., B.S.B., L.C., R.D.T., W.R.K., D.R., and M.S.; formal analysis and software, K.U., N.S., S.S.C., M.N., K.Y., and M.S.; resources, N.M. and N.S.; writing—original draft, M.S.; writing—review and editing, all authors. Supervision, G.L., U.G.S, J.M.G., and M.S.

## Corresponding authors

Correspondence to Masako Suzuki.

