## Supplemental figures 1-7 for "Prenatal vitamin D deficiency alters immune cell proportions of young adult offspring through alteration of long-term stem cell fates"

#### Table of Contents

**Supplementary Fig. 1:** The effects of vitamin D deficient diet feeding on the mothers and the impacts of prenatal VDD on the growth and bone density of offspring at the adult stage.

**Supplementary Fig. 2:** Prenatal VDD effects on female offspring at the adult stage.

**Supplementary Fig. 3:** Gating and analytical strategies to assess the immune cell profiles in peripheral blood and spleen.

**Supplementary Fig. 4:** Prenatal vitamin D deficiency alters hematopoietic cell cellular compositions of bone marrow at the adult stage.

**Supplementary Fig. 5:** Gating and analytical strategies to assess hematopoietic stem cells, multipotent progenitor cells, and progenitor cells from bone marrow.

**Supplementary Fig. 6:** Differentially expressed genes (DEGs) in VDD MPP4 are enriched in DEGs of hematopoietic transcription factor knockout models.

**Supplementary Fig. 7:** The heatmaps indicate the association between known covariates and immune cell proportions.

### Supplementary Fig. 1

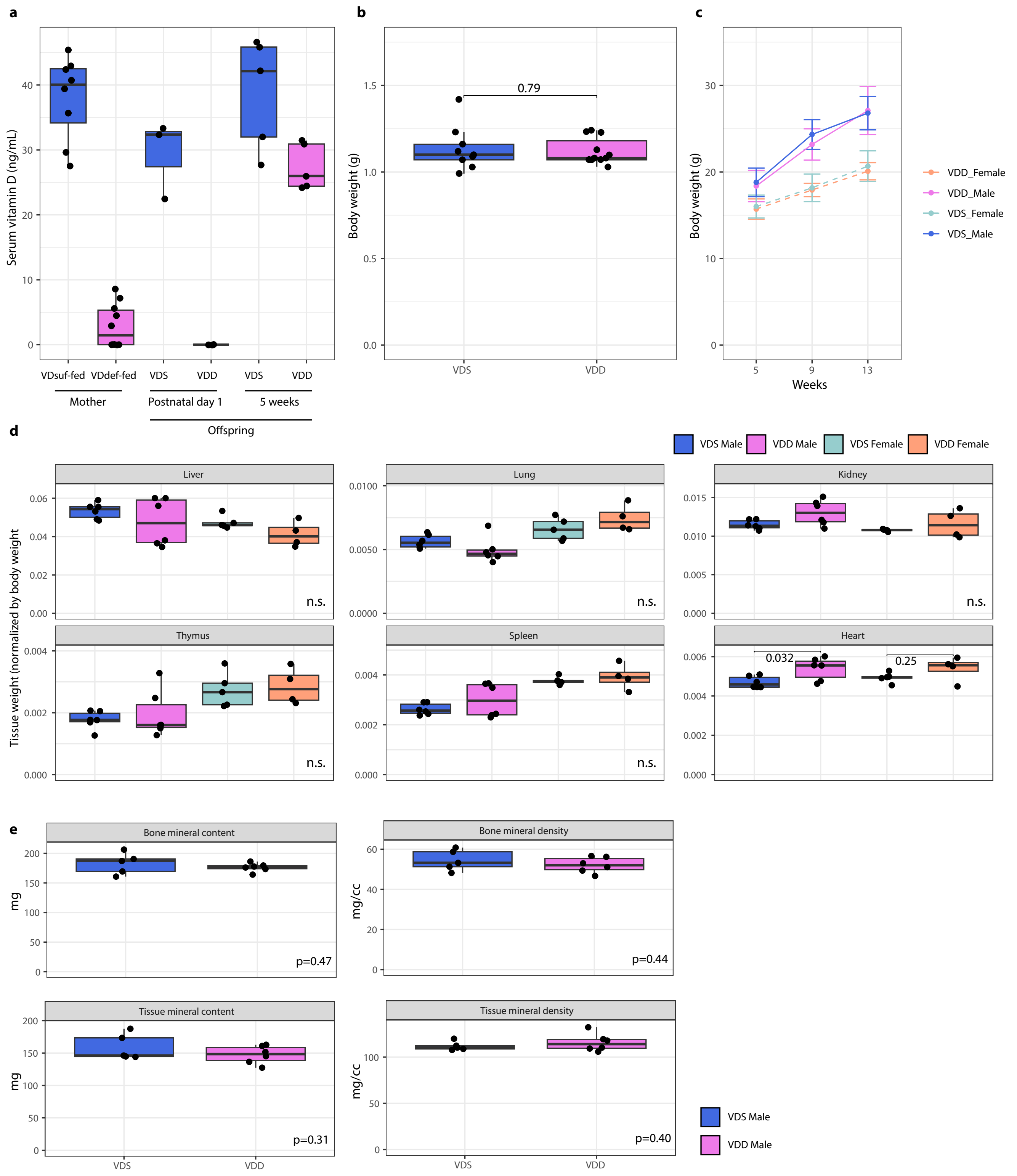

**Supplementary Fig. 1:** The effects of vitamin D deficient diet feeding on the mothers and the impacts of prenatal vitamin D deficiency on the growth and bone density of offspring at the adult stage.

**a** The deficient status of VDD mothers was confirmed by the serum vitamin D level of female mice (n=8 VDD mothers and n=10 VDS mothers) after five weeks of vitamin D-deficient diet feeding before the mating. The deficient status was maintained in the offspring at postnatal day 1 (n=3 per group). After weaning to the vitamin D sufficient diet, the vitamin D status of VDD offspring becomes sufficient at 5 weeks old (n=5 per group). **b** The body weight at postnatal day 1 was comparable between VDD and VDS offspring (n=9 VDS and n=11 VDD). **c** The growth was also comparable between VDD and VDS offspring in both males (n=28 VDD and n=15 VDS) and females (n=12 VDD and n=12 VDS). **d** The weight of each tissue was comparable in VDD (6 males and 4 females) and VDS (6 males and 5 females), except for the heart in males. **e** The micro CT analyses show that prenatal vitamin D deficiency does not affect the offspring's bone and tissue mineral status in adults (n=5 VDD males and n=6 VDS males).

**Supplementary Fig. 2**

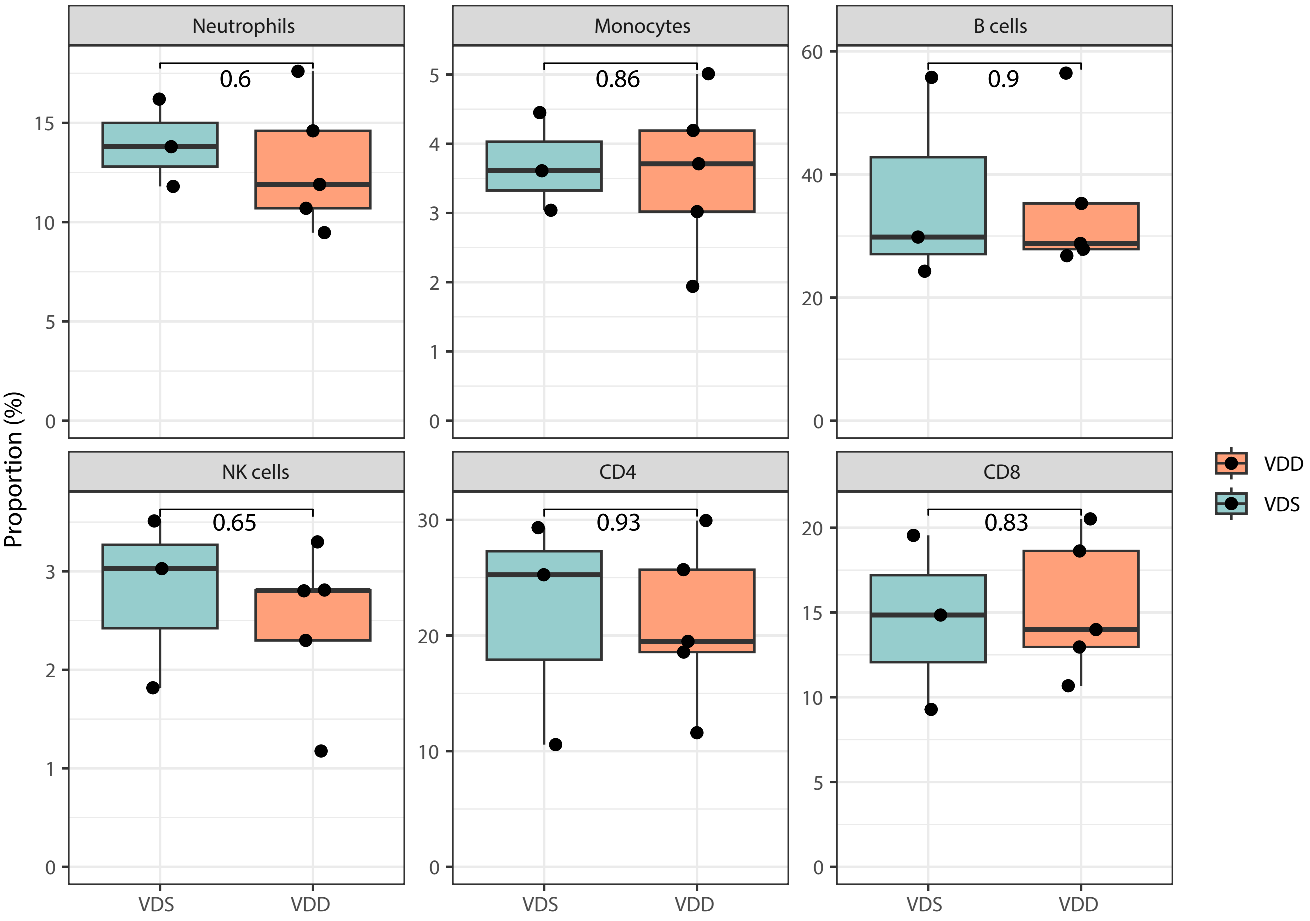


**Supplementary Fig. 2:** Prenatal vitamin D deficiency doesn't alter the immune cell proportions of female offspring at the adult stage.

The violin plots illustrate the proportions of immune cells in the peripheral blood of female mice at the adult stage, showing no significant alterations in the female samples (n=5 VDD and n=3 VDS). The white box shows the range between the first and third quartiles. The upper and lower whiskers represent the 1.5x inter-quantile range, while the black bars show the median. The values in the plot are p-values (Student's t-test).

**Supplementary Fig. 3**

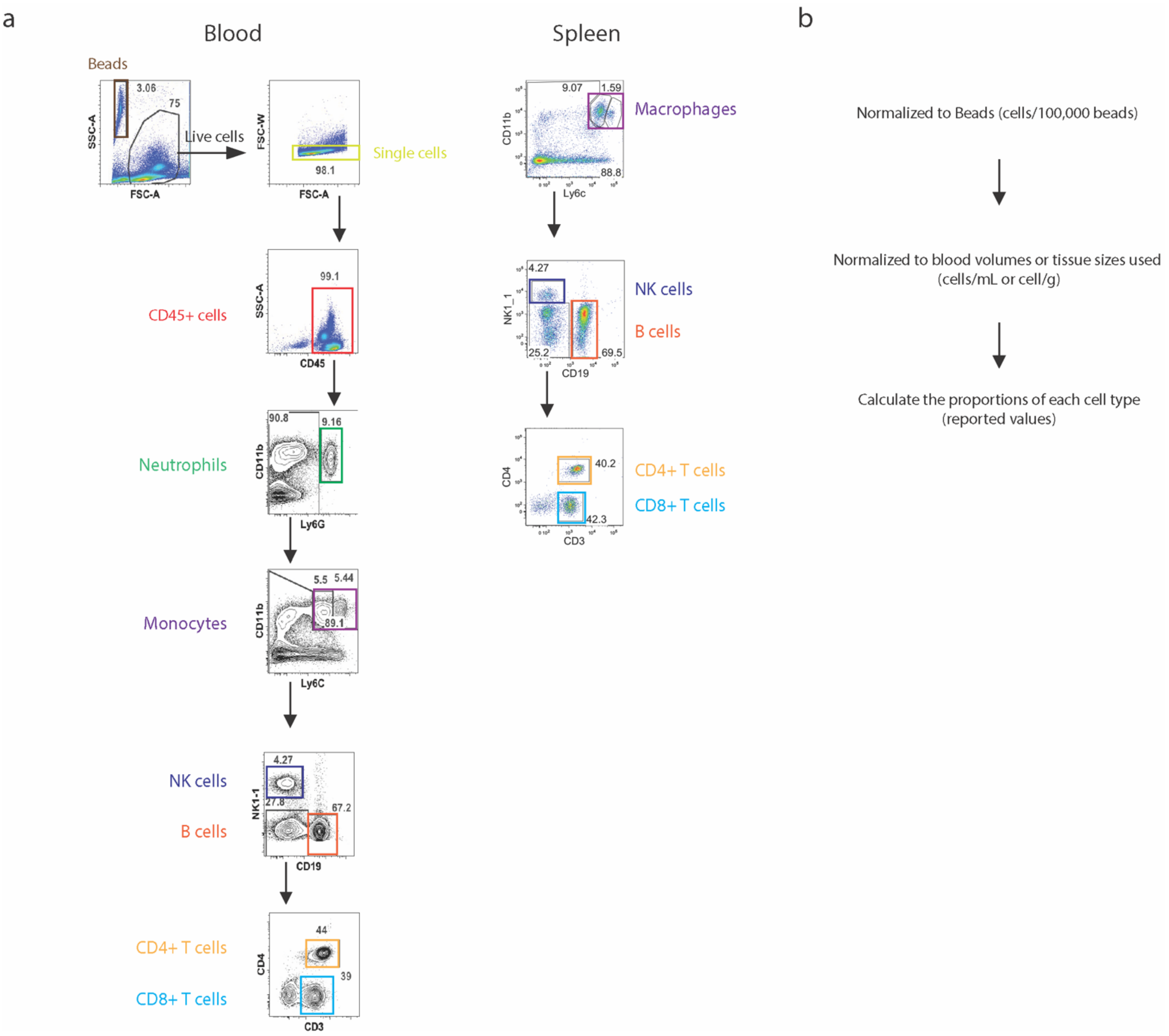


**Supplementary Fig. 3:** Gating and analytical strategies to assess the immune cell profiles in peripheral blood and spleen.

Representative FACS traces show the gating strategies to assess peripheral blood (**a**) and spleen (**b**). **c**. A flowchart outlines the process for calculating cell proportions. The panels on the left display the results from the VDS male mouse, while the panels on the right show the results from the VDD male mouse. The cell numbers obtained from the FACS results were first normalized by the number of beads we spiked into the samples before the wash steps and then further normalized by the blood volumes or tissue weight (spleen). The obtained cell numbers were used to calculate immune cell proportions.

#### Supplementary Fig. 4

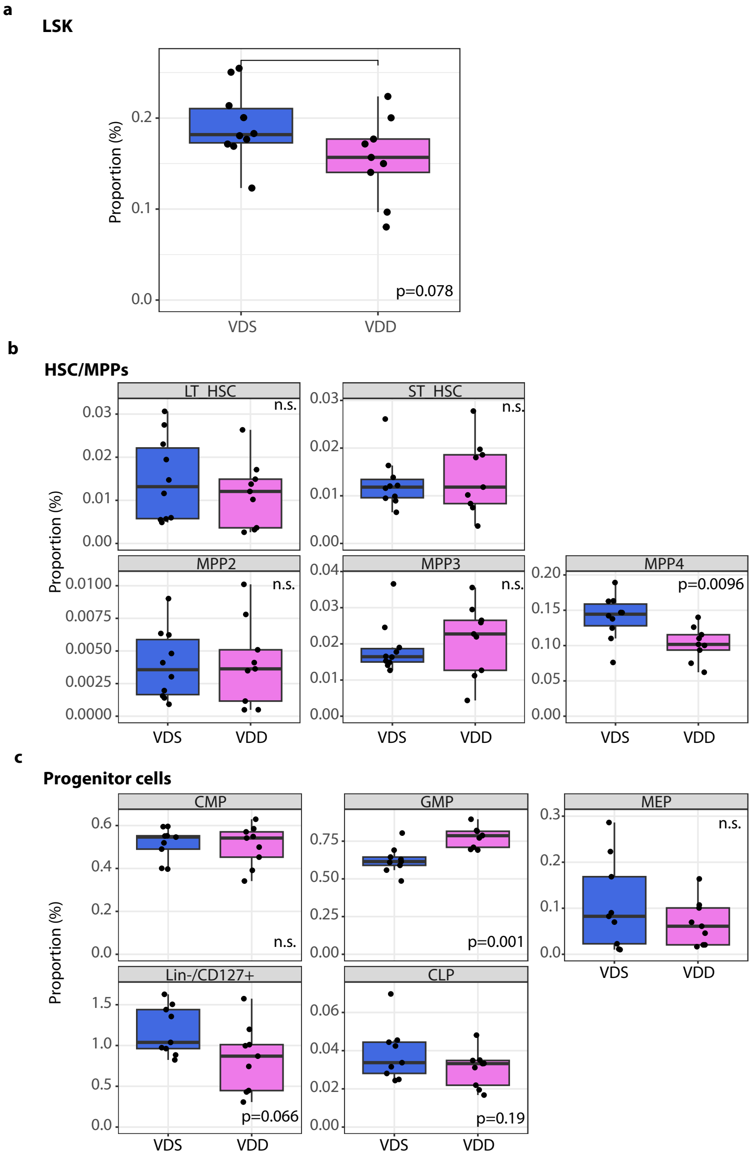


**Supplementary Fig. 4: Prenatal vitamin D deficiency alters hematopoietic cell cellular compositions of bone marrow at the adult stage.**

(**a**) The percentage of LSK fraction in bone marrow cells is reduced in VDD mice (n=9 for VDD and n=10 for VDS). (**b**) Variations in the percentages of total HSC, long-term and short-term HSC, and three MPPs in the bone marrow of VDD and VDS mice are presented in the box plots (n=9 for VDD and n=10 for VDS). The proportion of MPP4 was significantly reduced in VDD (p=0.0096). (**c**) The box plots show the percentages of hematopoietic progenitor cells in the bone marrow (n=9 per group). The proportion of GMP significantly increased in VDD (p=0.001).
Each dot on the plot represents a sample. The box represents the range between the first and third quartiles. The upper and lower whiskers represent 1.5 times the interquartile range, while the black bars indicate the median. The values displayed in the plot are p-values calculated using Student's t-test.

#### Supplementary Fig. 5

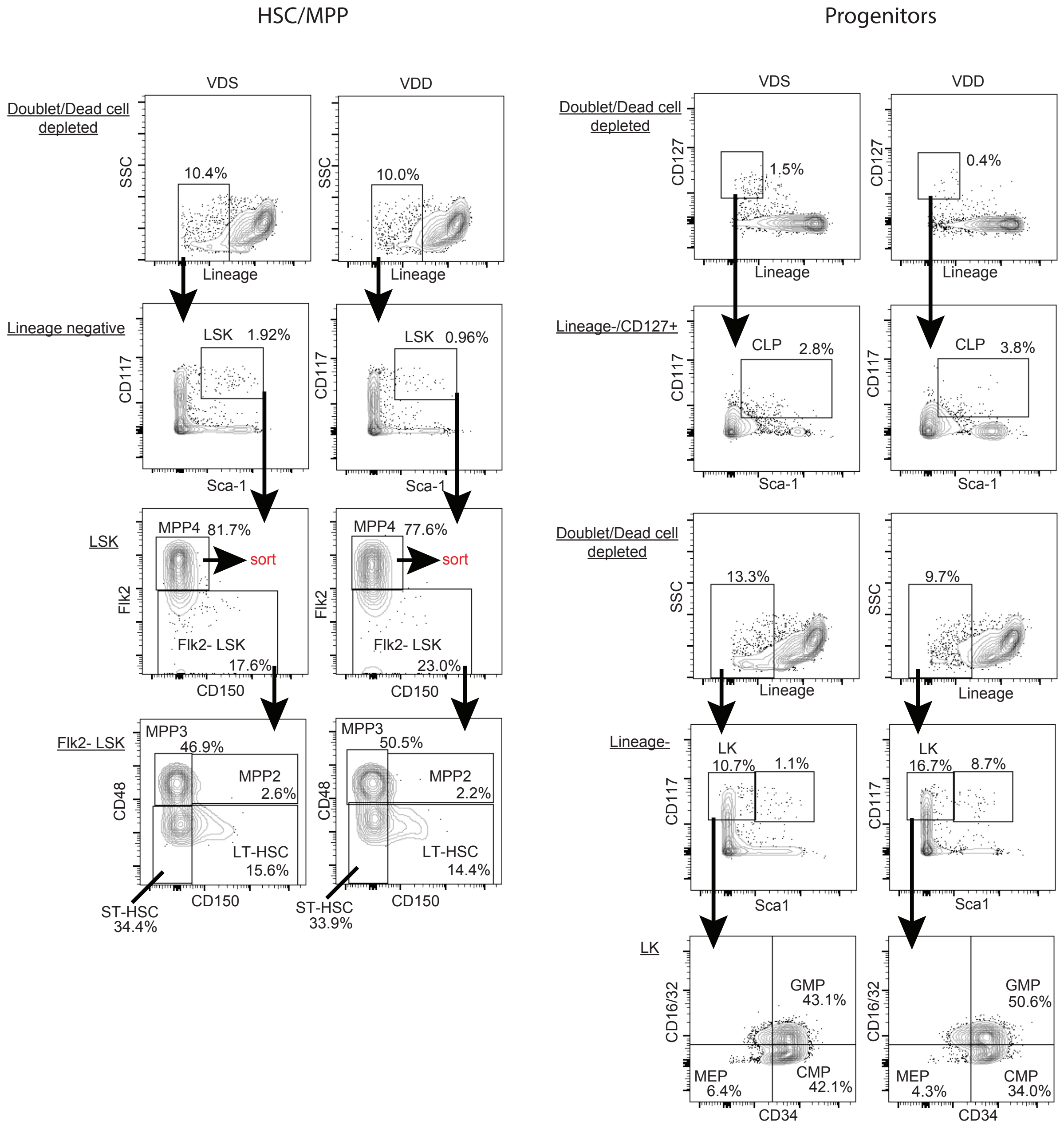


**Supplementary Fig. 5:** Gating and analytical strategies to assess hematopoietic stem cells, multipotent progenitor cells, and progenitor cells from bone marrow.

The gating strategies used to assess hematopoietic stem cells (HSC) and multipotent progenitor cells (MPPs) (a) and progenitor cells (b) are shown in the representative FACS traces. The results from the VDS male mouse are displayed in the panels on the left of the flow chart, while the panels on the right show the results from the VDD male mouse. The sorted cell populations used for RNA-seq analysis are indicated as "sort" in the bottom second panels of the HSC/MPP flow chart.

**Supplementary Fig. 6**

**Upregulated genes**

| | Term | Overlap | Adj P-value | Odds Ratio |
|---|---|---|---|---|
| * | NFIX KD MOUSE GSE45492 CREEDSID GENE 2846 UP | 61/281 | 3.92E-47 | 19.82398061 |
| | CEBPA KO MOUSE GSE61468 CREEDSID GENE 1476 DOWN | 52/278 | 1.98E-36 | 15.91868485 |
| | POU2AF1 OE MOUSE GSE12421 CREEDSID GENE 2312 DOWN | 50/261 | 1.25E-35 | 16.29125986 |
| | E2F8 KO MOUSE GSE52157 CREEDSID GENE 2828 DOWN | 58/406 | 1.79E-34 | 11.70848485 |
| $ | SRF KO MOUSE GSE23556 CREEDSID GENE 175 UP | 43/246 | 9.15E-29 | 14.21695261 |
| | MBD2 KO MOUSE GSE48653 CREEDSID GENE 1051 UP | 48/389 | 2.62E-25 | 9.545176725 |
| | HMGA2 OE MOUSE GSE24071 CREEDSID GENE 2564 DOWN | 42/289 | 7.70E-25 | 11.34771902 |
| | DNMT1 KO MOUSE GSE31626 CREEDSID GENE 1236 DOWN | 43/325 | 7.84E-24 | 10.19265101 |
| | BATF KO MOUSE GSE54215 CREEDSID GENE 1522 UP | 36/215 | 2.36E-23 | 13.19654647 |
| | DNMT1 KO MOUSE GSE31626 CREEDSID GENE 1237 DOWN | 38/292 | 1.06E-20 | 9.845108768 |

**Downregulated genes**

| | Term | Overlap | Adj P-value | Odds Ratio |
|---|---|---|---|---|
| | RUNX1 KO MOUSE GSE40155 CREEDSID GENE 1492 UP | 39/286 | 8.91E-25 | 13.55460236 |
| * | NFIX KD MOUSE GSE45492 CREEDSID GENE 2846 DOWN | 30/188 | 8.00E-21 | 15.74983909 |
| | CEBPA KO MOUSE GSE61468 CREEDSID GENE 1476 UP | 31/238 | 4.89E-19 | 12.44397163 |
| $ | SRF KO MOUSE GSE23556 CREEDSID GENE 175 DOWN | 25/258 | 4.55E-12 | 8.682082881 |
| | ZBTB7A KO MOUSE GSE41839 CREEDSID GENE 2380 DOWN | 23/246 | 9.53E-11 | 8.281256344 |
| | NFE2L2 KO MOUSE GSE18344 CREEDSID GENE 1259 DOWN | 28/419 | 7.72E-10 | 5.820069204 |
| | KLF13 KO MOUSE GSE25502 CREEDSID GENE 1821 UP | 21/290 | 9.52E-08 | 6.202336697 |
| | NFE2L2 KO MOUSE GSE18344 CREEDSID GENE 1258 DOWN | 23/406 | 1.39E-06 | 4.782183111 |
| | ZFX KO MOUSE GSE7069 CREEDSID GENE 178 UP | 16/206 | 3.29E-06 | 6.583242105 |
| | GATA3 KO MOUSE GSE39864 CREEDSID GENE 1162 DOWN | 16/206 | 3.29E-06 | 6.583242105 |

**Supplementary Fig. 6:** Differentially expressed genes (DEGs) in VDD MPP4 are enriched in DEGs of hematopoietic transcription factor knockout models.

The bar charts show the enrichments of DEGs of VDD MPP4 in the DEGs of the top 10 enriched transcription factor knock-down, knockout, or over-expression models (sorted by p-values). **Top** Upregulated genes in VDD MPP4. **Bottom** Downregulated genes in VDD MPP4. * and # indicate that the enrichments were observed in both up- and down-regulated genes.

**Supplementary Fig. 7**

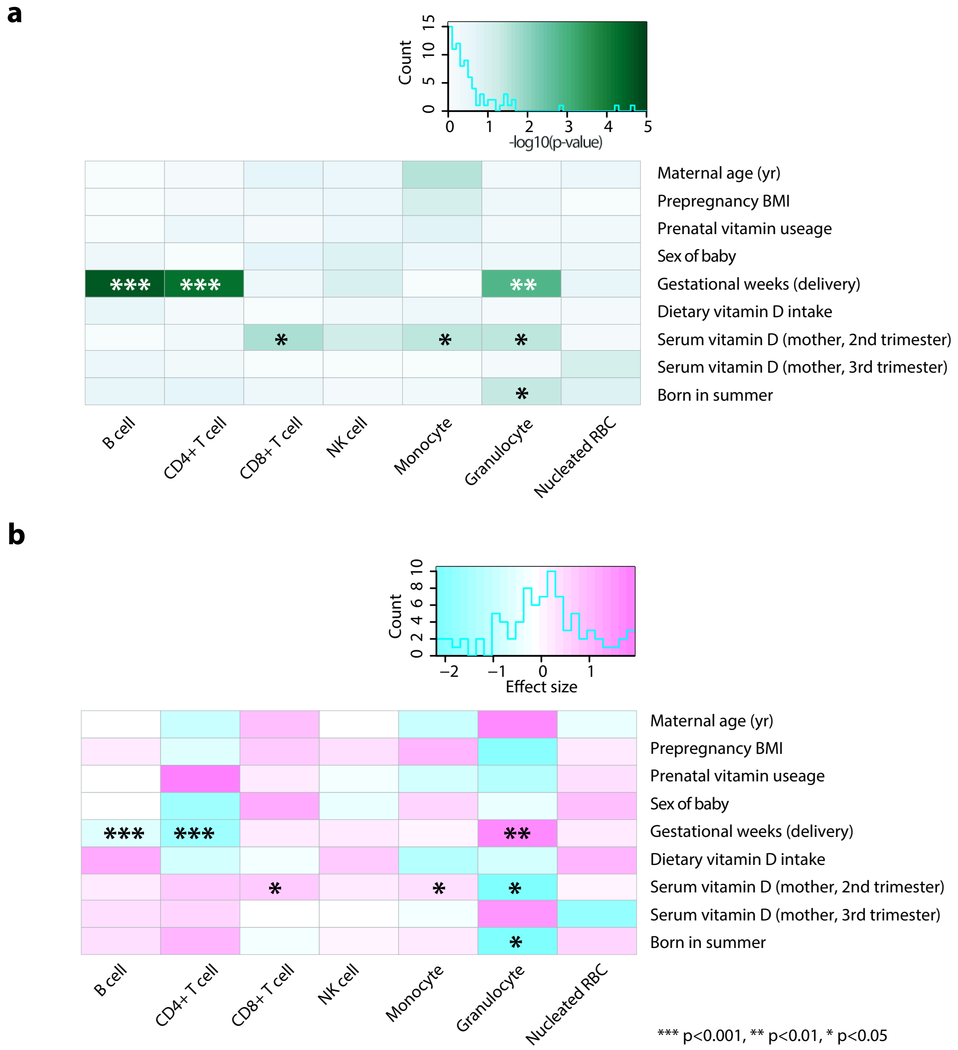


**Supplementary Fig. 7:** The association between known covariates and immune cell proportions of cord blood

The heatmaps show that the gestational week at the delivery, maternal serum vitamin D (2$^{nd}$ trimester), and being born in the summer season are significantly associated with the proportions of immune cells. The heatmaps represent -log10(p-value) (**a**) and the direction of the associations (**b**). Asterisks indicate the significance (*** p<0.001, ** p<0.01 and * p<0.05, Student's t-test).
